# Bypassing PELO-mediated ATPase activation of NLR is a common pathogenic cause of *NLR*-associated autoinflammatory diseases

**DOI:** 10.1101/2024.01.20.576395

**Authors:** Xiurong Wu, Zhang-Hua Yang, Yue Zheng, Jianfeng Wu, Jiahuai Han

## Abstract

Nucleotide-binding and oligomerization domain (NOD)-like receptors (NLRs) constitute the largest family of pattern recognition receptors. These receptors are master regulators of innate immunity. Recently, PELO-driven ATPase activation of NLRs was demonstrated as a critical step in NLR activation. Linkage studies in human with various inherited autoinflammatory conditions have revealed an extensive array of mutations and polymorphisms within NLRs. However, the precise mechanisms by which genetic variations in NLR genes contribute to disease onset remain largely elusive. Here we comprehensively analyze dozens of naturally occurring mutations across multiple human NLRs, and demonstrate that *NLRs* harboring pathogenic mutations in their NACHT domain—not those with non-pathogenic variants—exhibit spontaneous PELO-independent ATPase activation. This aberrant activation triggers corresponding NLR to mediate inflammatory responses. Thus, bypassing the PELO-checkpoint for ATPase activation is a major disease-causing mechanism underlying *NLR* mutations. Furthermore, quantifying this aberrant ATPase activation could serve as an assessment tool for classifying the pathogenesis of *NLR*-associated diseases.

## INTRODUCTION

Nucleotide-binding and oligomerization domain (NOD)-like receptors (NLRs) are evolutionarily conserved intracellular pattern recognition receptors (PRRs) that translate microbial and danger sensing into immediate host defenses and prime the adaptive immune response for long-lasting protection^1^. In humans, 22 known NLRs exist, and mutations in NLR family genes are associated with a wide range of inflammatory and autoimmune conditions, including hereditary periodic fever syndromes, Crohn ‘s disease, Blau ‘s syndrome, infantile enterocolitis, multiple sclerosis, and asthma^2,3^. Collectively, these disorders can be categorized as *NLR*-associated autoinflammatory diseases.

The hallmark feature of NLRs is their central NOD (or NACHT) domain, while the N-terminal and C-terminal domains exhibit variability. These domains may include homotypic protein-protein interaction domains and leucine-rich repeats (LRRs)^4^. The NACHT domain—an ATPase domain, comprises several subdomains: nucleotide binding domain (NBD), helical domain 1, winged-helix domain, and helical domain 2. Upon ligand binding, the auto-inhibitory C-terminal LRR domain of most NLRs undergoes a conformational change, exposing the N-terminal domain. This conformational shift allows for the formation of oligomeric scaffolds, which subsequently interact with downstream signaling adaptors or effectors. Notably, ATPase activity embedded within NLRs is essential for complex formation^4^. Specific NLRs, such as NLRP1, NLRP3, NLRP6, NLRP7, NLRP12, NLRC4, and NAIP, operate *via* inflammasomes to elicit inflammatory responses. Conversely, NOD1, NOD2, NLRP10, NLRX1, NLRC5, and CIITA activate inflammatory signaling pathways, including nuclear factor-κB (NF-κB), mitogen-activated protein kinases (MAPKs), and interferon (IFN) regulatory factors (IRFs)^3^. Despite the diversity of inflammatory pathways activated by different NLR family members, recent research has revealed a common checkpoint of ATPase activation governing the activation of all NLRs^5^.

PELO (Protein pelota homolog) is a conserved component of the ribosome-associated quality control machinery. It functions as a surveillance factor in translational quality control and ribosome rescue^6-10^. Recently, we have a surprising discovery that PELO is essential for the activation of all cytosolic NLR family proteins. The underlying mechanism involves PELO recruitment by NLR proteins upon activation, catalyzing the activation of the ATPase within the NACHT domain of NLRs^5^. The effect(s) of the pathogenic mutations of NLRs on their ATPase activation by PELO deserves investigation.

## RESULTS

### Pathogenic NLRP3 mutations lead to bypassing PELO-mediated ATPase activation of NLRP3

To initiate our analysis, we focused on NLRP3, one of the most extensively studied NLR family members^11^. NLRP3 forms an inflammasome—an intracellular multimeric protein complex—in response to various exogenous microbial infections and endogenous danger signals^12^. Numerous mutations have been reported in NLRP3, and these mutations are associated with autoinflammatory diseases such as cryopyrin-associated periodic syndromes (CAPS). CAPS includes familial cold autoinflammatory syndrome (FCAS), Muckle-Wells syndrome (MWS), and chronic infantile neurological, cutaneous, and articular syndrome (CINCA, also known as NOMID)^13^. Interestingly, CAPS can occur not only in familial forms but also sporadically due to *de novo* or somatic mosaic NLRP3 mutations^14^. In our investigation, we initially analyzed the five most frequent gain-of-function germline mutants of NLRP3 to assess their impact on ATPase activation by PELO^14^. We expressed and purified NLRP3 and its mutants from *PELO* knockout HEK293T cells (**Figure S1A**) and measured their ATPase activities. Consistent with previous reports^5^, the addition of purified recombinant PELO protein substantially increased the ATPase activity of NLRP3 (**Figure 1A**). Interestingly, these five NLRP3 mutants already exhibited high ATPase activity, and the addition of PELO had minimal or no effect on further enhancing their ATPase activity (**Figure 1A**). Consequently, these disease-causing NLRP3 mutants appear capable of bypassing the PELO-mediated activation step and achieving self-activation.

**Figure 1.**
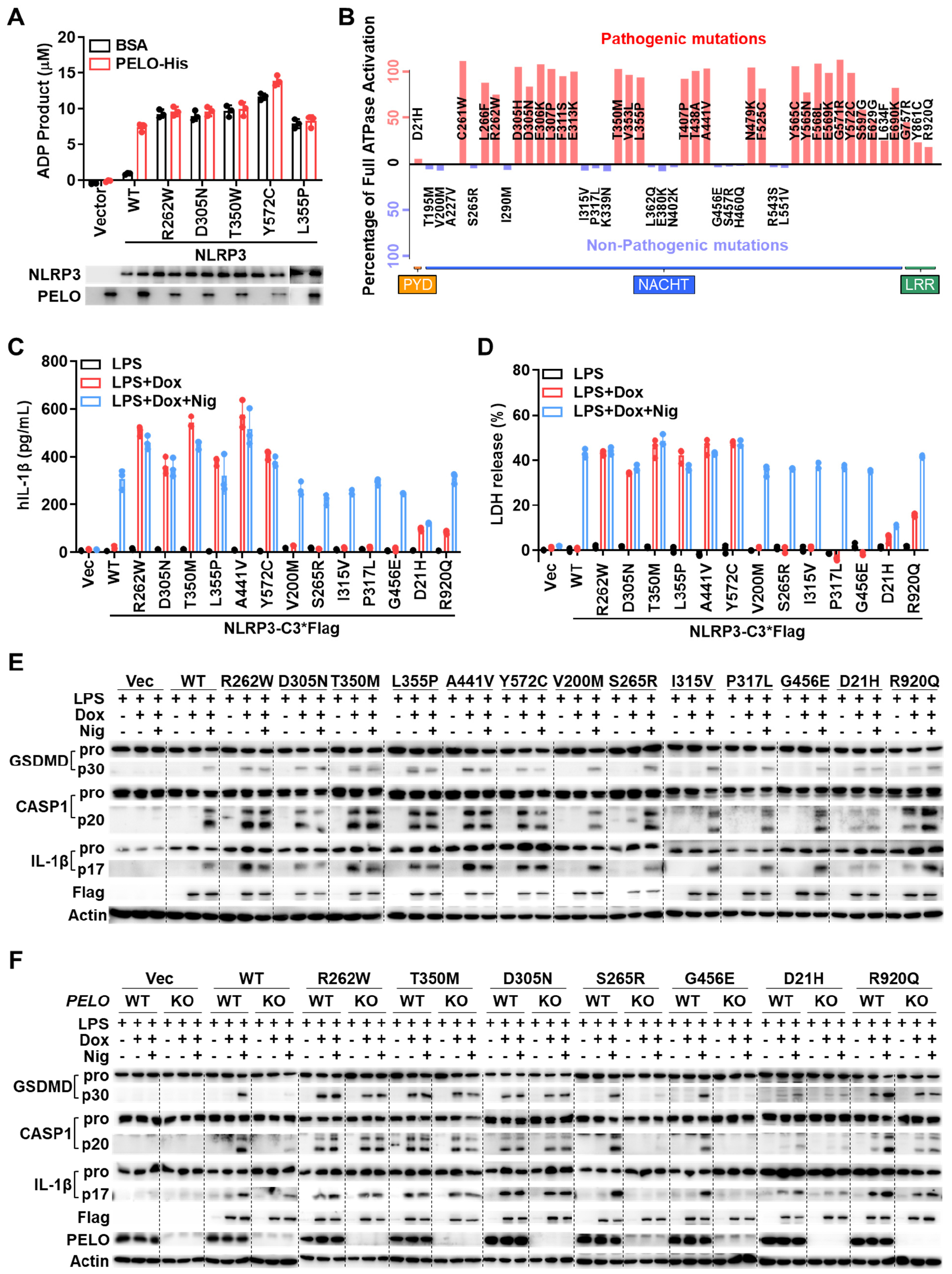
Pathogenic NLRP3 mutations lead to bypassing PELO-mediated ATPase activation of NLRP3 by gain of ATPase activity and each of the gain-in-function NLRP3 mutations spontaneously elicits inflammasome activation. (A) Purified Flag-WT or one of the five mutated NLRP3 proteins (100 ng) as indicated were incubated with BSA or recombinant PELO protein (50 ng) in the presence of 20 μM ATP, and ADP product was measured using the ADP-Glo assay. (B) The experiments were performed as in (**A**) with 47 NLRP3 mutants. Schematics show the basal level ATPase activity of each NLRP3 mutant relative to the full activation induced by incubation with PELO. Pathogenic mutations are in red and non-pathogenic mutations are in blue. The positions of the bars of each mutant in the schematic diagram correspond to the domain location of each mutation site. **(C-E)** *NLRP3* KO THP-1 cells with stable integration of Dox-inducible expression vector of Flag-NLRP3 WT or one of the selected NLRP3 mutants were firstly primed with 1 mg ml^-1^ LPS for 4 hours. Following priming, the medium was replaced with medium with or without 1 μg ml^-1^ doxycycline (Dox). Six hours later, the cells were stimulated with or without 5 μM nigericin (Nig) for another 1 hour. Supernatants were analyzed for IL-1β(C) and LDH (**D**); the pooled cell extracts and supernatants were analyzed by immunoblotting with anti-GSDMD, anti-caspase-1 (CASP1), anti-IL-1β and anti-actin (**E**). **(F)** *NLRP3* KO THP-1 cells with and without additional *PELO* KO were stably transfected with Dox-inducible expression vector of Flag-NLRP3 WT or one of the designated mutants, and were treated with the stimuli as in (**E**). The processed GSDMD, caspase-1 (CASP1) and IL-1β in the pooled cell extracts and supernatants were analyzed by immunoblotting. Equal loading of the proteins in ATPase activity assay was monitored by Western blotting. Data are represented as mean ± SD of triplicates (**A, C** and **D**). All results are representative of at least two independent experiments. See also **Figure S1**.

To investigate whether bypassing PELO-mediated ATPase activation is a widespread phenomenon among disease-causing NLRP3 mutants, we expanded our analysis to include 47 naturally occurring NLRP3 mutations. This comprehensive set encompasses germline and somatic mosaic mutants, as well as pathogenic and non-pathogenic variants^14^. **Figure 1B** provides insights into the domain locations of these mutants, their disease-causing potential, and illustrates the basal activity levels of each mutant relative to the full activation achieved through incubation with PELO. Their ATPase activities with and without PELO activation are shown in **Figure S1B**. Our findings reveal a consistent pattern: without exception, pathogenic mutations within the NACHT domain exhibit basal ATPase activation, whereas non-pathogenic mutants require PELO for ATPase activation (**Figures 1A, 1B and S1B**). Interestingly, mutations occurring outside the NACHT domain—such as those in the LRR domain—can also gain certain levels of ATPase activity, which PELO further enhances (**Figures 1B and S1B**). Remarkably, even these LRR mutations only exhibit modest basal ATPase activity they are pathogenic. It needs to note that the impact of NLRP3 mutations on PELO-mediated ATPase activation is independent of their ability to interact with PELO (**Figure S1C**). Thus, the ability to bypass the PELO checkpoint and directly self-activate ATPase emerges as a common feature among pathogenic NLRP3 mutants.

### Pathogenic NLRP3 mutations that bypass PELO-mediated ATPase activation spontaneously elicits inflammasome activation

To investigate whether gain-of-function mutations in ATPase activates NLRP3, we analyzed inflammasome activation using a *NLRP3* knockout human monocyte cell line (THP-1). Key features of inflammasome activation include caspase-1 activation, which processes proinflammatory cytokines interleukin-1β (IL-1β) and/or IL-18 for maturation, and cleavage of gasdermin D (GSDMD) to generate the N-terminal fragment, inducing pore formation, cytokine release, and pyroptosis^11^. While in standard cell model NLRP3 and pro-IL-1β expression is typically induced by lipopolysaccharide (LPS), our study employed *NLRP3* knockout THP-1 cells with the expression of NLRP3 or its mutants achieved *via* a doxycycline (Dox)-inducible system. Thirteen mutants, harboring both pathogenic and non-pathogenic mutations across distinct domains of NLRP3, were selected for evaluation. As in the standard experimental procedure, nigericin (Nig) was employed to induce caspase-1 activation, IL-1β maturation and secretion, and pyroptosis in LPS-primed THP-1 cells. The experiment procedure was: following LPS priming, cells were stimulated with either nothing, Dox alone, or Dox in combination with Nig. IL-1β secretion was quantified by assessing IL-1β levels in the cell culture medium (**Figure 1C**), while pyroptosis was evaluated through lactate dehydrogenase (LDH) release into the medium (**Figure 1D**). Additionally, IL-1β maturation, caspase-1 cleavage, and gasdermin D (GSDMD) cleavage were determined by Western blotting (**Figure 1E**). Results revealed an unresponsive phenotype in *NLRP3* knockout THP-1 cells to LPS, LPS+Dox, and LPS+Dox+Nig. Induction of NLRP3 expression by Dox reinstated responses to LPS+Nig, bringing them to a level comparable to wildtype THP-1 cells. The induced expression of NLRP3 mutants with spontaneous ATPase activation directly triggers caspase-1 activation, GSDMD cleavage, IL-1β maturation and secretion in the absence of secondary stimulation (nigericin). In contrast, mutants that are not gain-of-function of ATPase activity do not directly activate the NLRP3 inflammasome; instead, they behave similarly to wildtype NLRP3 and respond to nigericin. Notably, the D21H mutant exhibits distinct behavior from other pathogenic mutants, likely due to its location in the PYD domain, which interacts with downstream adaptor protein ASC^15^. Collectively, these data support the notion that ATPase activation by PELO serves as a checkpoint for NLRP3 inflammasome activation, and the gain-of-function ATPase mutations bypass this checkpoint.

To confirm that PELO controls this checkpoint, we conducted the aforementioned experiments in the cells with and without additional *PELO* deletion. As expected, *PELO* deficiency only affected wildtype NLRP3 and the mutants that did not gain basal ATPase activity (**Figure 1F**). Conclusively, each gain-of-function ATPase mutation in NLRP3 bypassed PELO for ATPase activation, resulting in spontaneous inflammasome activation.

### Bypass of PELO-mediated ATPase activation is a common mechanism underlying the pathogenicity of disease-causing mutations of NLR family members

Next, we investigated whether the conclusions drawn from the study of NLRP3 could be extended to other NLR family members. NLRC4, another well-studied NLR, is associated with autoinflammatory diseases such as autoinflammation and infantile enterocolitis (AIFEC)^16^. Similar to our NLRP3 study, we selected specific mutations in NLRC4 and assessed their ATPase activation by PELO *in vitro* (**Figures 2A and S2A**). The results mirrored those obtained for NLRP3: all pathogenic mutations (highlighted in red) in the NACHT domain bypassed the PELO checkpoint and gained basal ATPase activity. Conversely, PELO remained necessary for ATPase activation in non-pathogenic mutations (highlighted in blue) within the NACHT domain. In line with the NLRP3 findings, pathogenic mutations in the CARD domain showed no functional relationship with the PELO checkpoint for ATPase activation, while LRR domain mutations partially bypassed this checkpoint. Although mutations have been identified in most, if not all, NLR family members, disease-associated studies have primarily focused on NLRP3 and NLRC4 with additional NOD1, NOD2, NLRP7, and NLRP12. To further explore the impact of PELO-checkpoint on these additional NLRs, we examined PELO ‘s effect on the ATPase activity of the pathogenic mutations in the NACHT domain of these NLRs. As depicted in **Figure 2B** and **Figures S2B-S2D**, all of these mutations led to the bypass of PELO-mediated ATPase activation. Our data collectively demonstrate that bypassing the PELO checkpoint through gain-of-function ATPase mutations represents a common pathogenic mechanism in *NLR*-associated autoinflammatory diseases.

**Figure 2.**
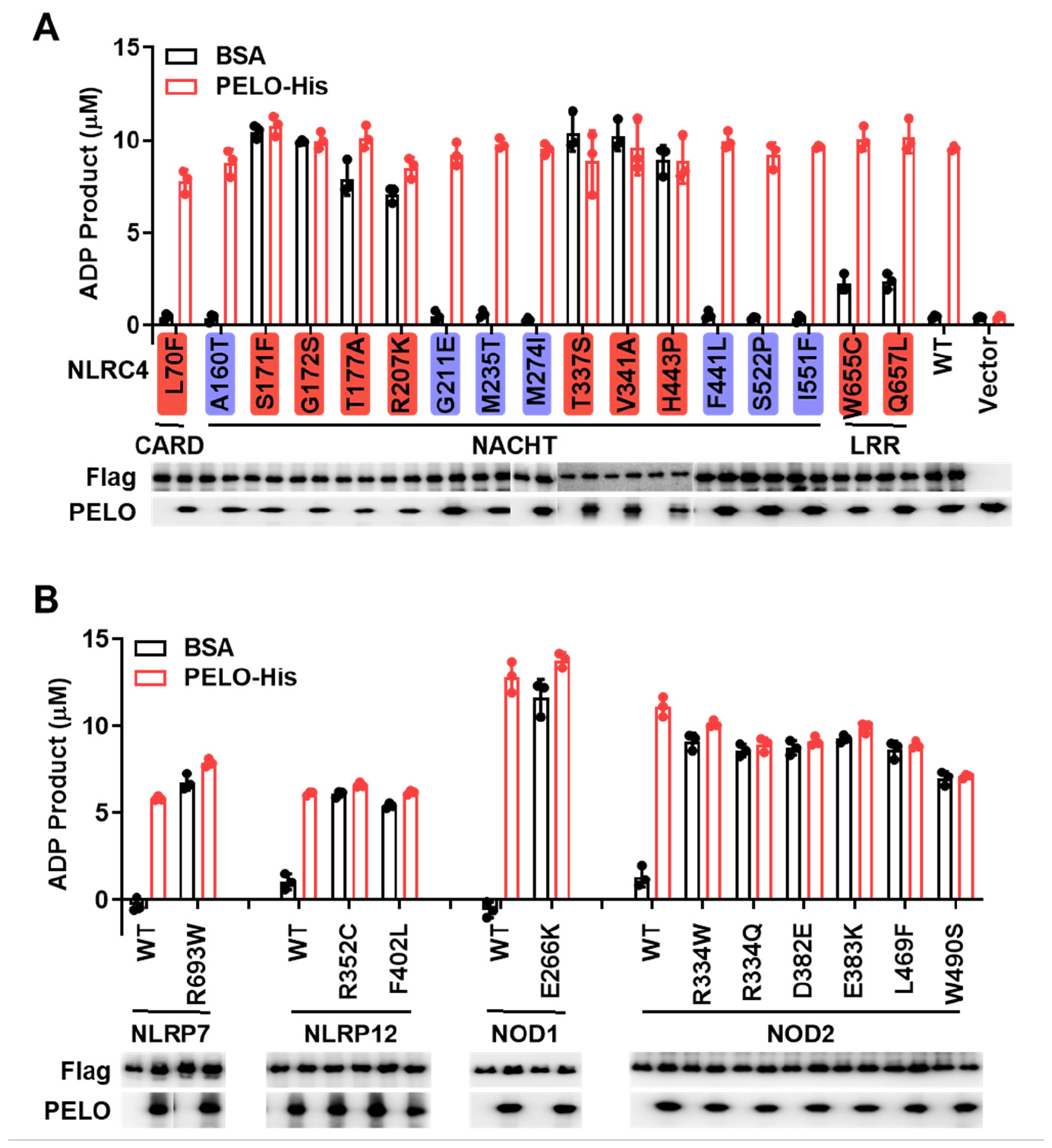
Bypass of PELO-mediated ATPase activation is a common mechanism underlying the pathogenicity of disease-causing mutations of NLR family members. **(A)** Purified proteins of Flag-WT NLRC4 or one of the selected NLRC4 mutants (100 ng) were incubated with BSA or recombinant PELO protein (50 ng) in the presence of 20 μM ATP, and ADP product was measured using the ADP-Glo assay. The pathogenic mutations were highlighted in red, and the non-pathogenic mutations were in blue. **(B)** Purified proteins of Flag-NLRP7, Flag-NLRP12, Flag-NOD1, Flag-NOD2 or their pathogenic mutations (100 ng) were incubated with BSA or recombinant PELO protein (50 ng) in the presence of 20 μM ATP, and ADP product was measured using the ADP-Glo assay. Equal loading of the proteins in ATPase activity assay was monitored by Western blotting. Data are represented as mean ± SD of triplicates. All results are representative of at least two independent experiments. See also **Figure S2**.

## DISCUSSION

Through the analysis of over 70 naturally occurring mutations across six human NLR family members, we unveil a common underlying mechanism driving the pathogenicity of NLR mutations: the bypass of the PELO checkpoint for ATPase activation in NLRs. Since the gain of ATPase activity by mutation is sufficient to trigger downstream inflammatory reactions of the respective NLRs., bypass of PELO-mediated ATPase activation is the pathogenic cause of the mjarity *NLR*-associated autoinflammatory diseases.

Among all the pathogenic mutations in the NACHT domain that we tested, the PELO checkpoint is consistently bypassed. However, beyond the NACHT domain, some mutations also circumvent the ATPase activation checkpoint. Thus, alternative pathogenic activation mechanisms exist. One such case involves the PYD domain mutant D21H. Although its ATPase activity can be activated by PELO, its inflammasome activation remains unresponsive to PELO, as demonstrated by the lack of response to nigericin (**Figure 1C-1E and S1B**). A plausible interpretation for this phenomenon is that the D21H mutation alters the structure of the PYD domain, enhancing its interaction with the adaptor protein ASC^15^. Consequently, it exhibits a certain level of low basal activity that remains unaffected by conformational changes in the NACHT domain.

As the identification of sequence variations (including germline and somatic mutations) in NLRs continues to grow, assessing whether the PELO checkpoint for ATPase activation has been bypassed can serve as a valuable tool for classifying pathogenesis in patients with NLR mutations. Computational modeling of NLRP3 has been instrumental in evaluating the potential relationship between clinical severity and the structural disruptions caused by mutations^17^, but its feasibility is not optimistic. By quantitatively determining both the extent of the bypass of ATPase activation checkpoint by a mutation and whether PELO can further enhance ATPase activity in a given NLR mutant, we can explore the clinical implications. This includes assessing vulnerability to sterile or infectious inflammation and the potential for excessive immune reactions.

Activation of the ATPase by PELO is an essential step in NLR activation, but how the PELO is regulated to participate this process is unknown. As for the case of NLRP3 activation, PELO ‘s participation is triggered by the second signal (nigericin), and the mechanism of PELO-checking point still awaits further investigation. Given the critical roles of NLRs in surveilling intracellular microbes, noxious substances, and other stress perturbations, the bypass of the PELO-mediated ATPase activation by mutations likely contributes to not only the NLR-associated autoinflammatory diseases but also numerous other adverse health conditions in humans.

## ACKNOWLEDGEMENTS

We thank Lu Zhou for help with proofreading. This work was supported by the National Natural Science Foundation of China (82388201 to J.H., 32170751 to Z.-H.Y.), National Key R&D Program of China (2020YFA0803500 to J.H.), the CAMS Innovation Fund for Medical Science (2019-I2M-5-062 to J.H.), Fujian province central to local science and technology development special program (No. 2022L3079) and Fu-Xia-Quan Zi-Chuang district cooperation program (No. 3502ZCQXT2022003).

## AUTHOR CONTRIBUTIONS

X.W. and Z.-H.Y. performed most of the experiments; Y.Z. and J.W. participated in the experiments; X.W., Z.-H.Y., and J. H. analyzed data and wrote the manuscript. X.W. and J.H. conceived the project and supervised the study.

## DECLARATION OF INTERESTS

The authors declare no competing interests.

**Figure S1.**
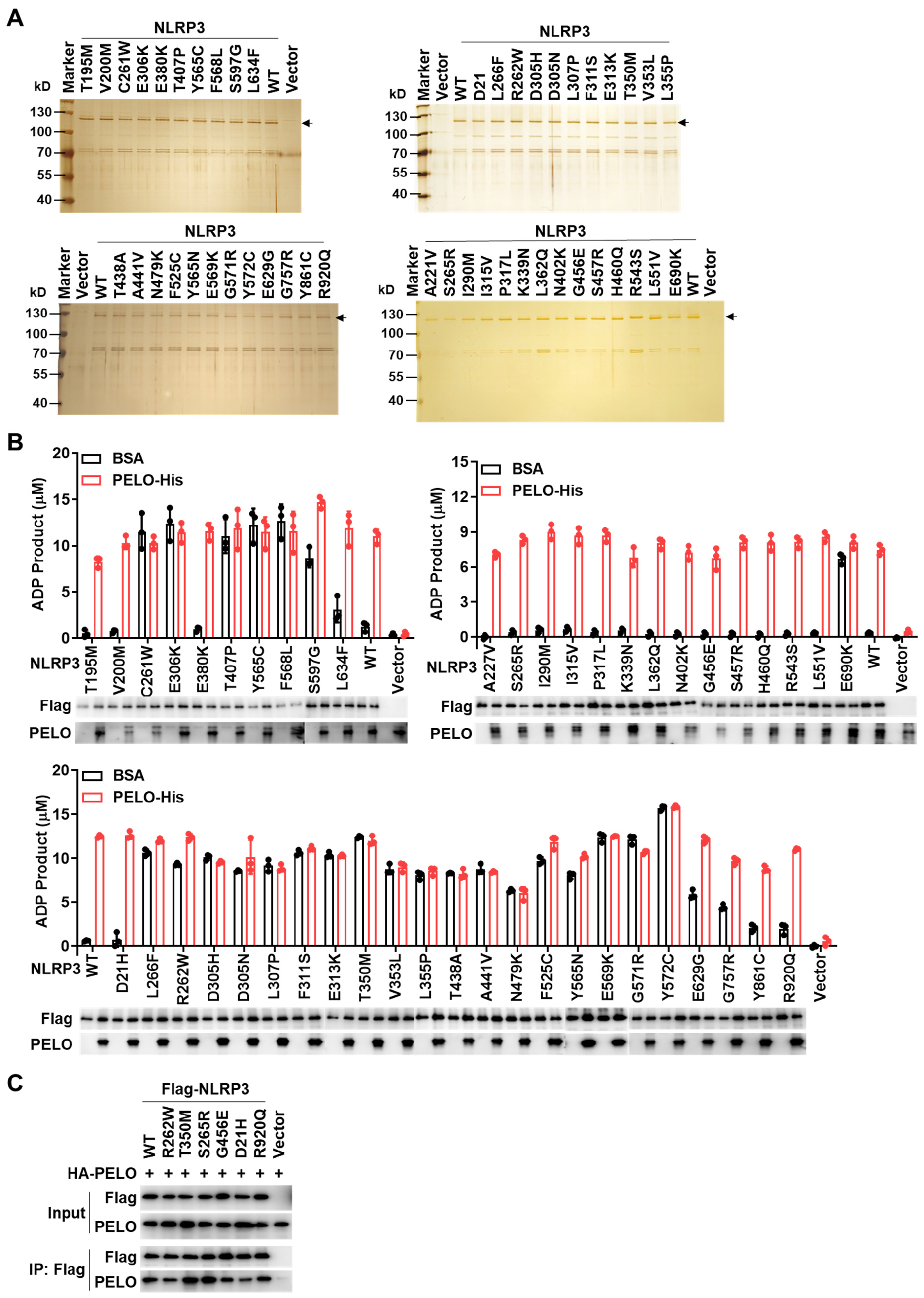
Purified proteins and ATPase activity assay, related to Figure 1. (A) Representative image of silver staining of the purified proteins of Flag-tagged NLRP3 and its mutants as indicated. (B) Purified Flag-WT or mutant NLRP3 proteins (100 ng) were incubated with BSA or recombinant PELO protein (50 ng) in the presence of 20 μM ATP, and the ADP product was measured using the ADP-Glo assay. Raw data of **Figure 1B**. (C) Flag-tagged WT or one of NLRP3 mutants as indicated was co-expressed with HA-tagged PELO in HEK293T cells. The cell lysates were immunoprecipitated with anti-Flag antibodies. The cell lysates and immunoprecipitates were analyzed by immunoblotting as indicated. Equal loading of the proteins in ATPase activity assay was monitored by Western blotting. Data are represented as mean ± SD of triplicates. All results are representative of at least two independent experiments.

**Figure S2.**
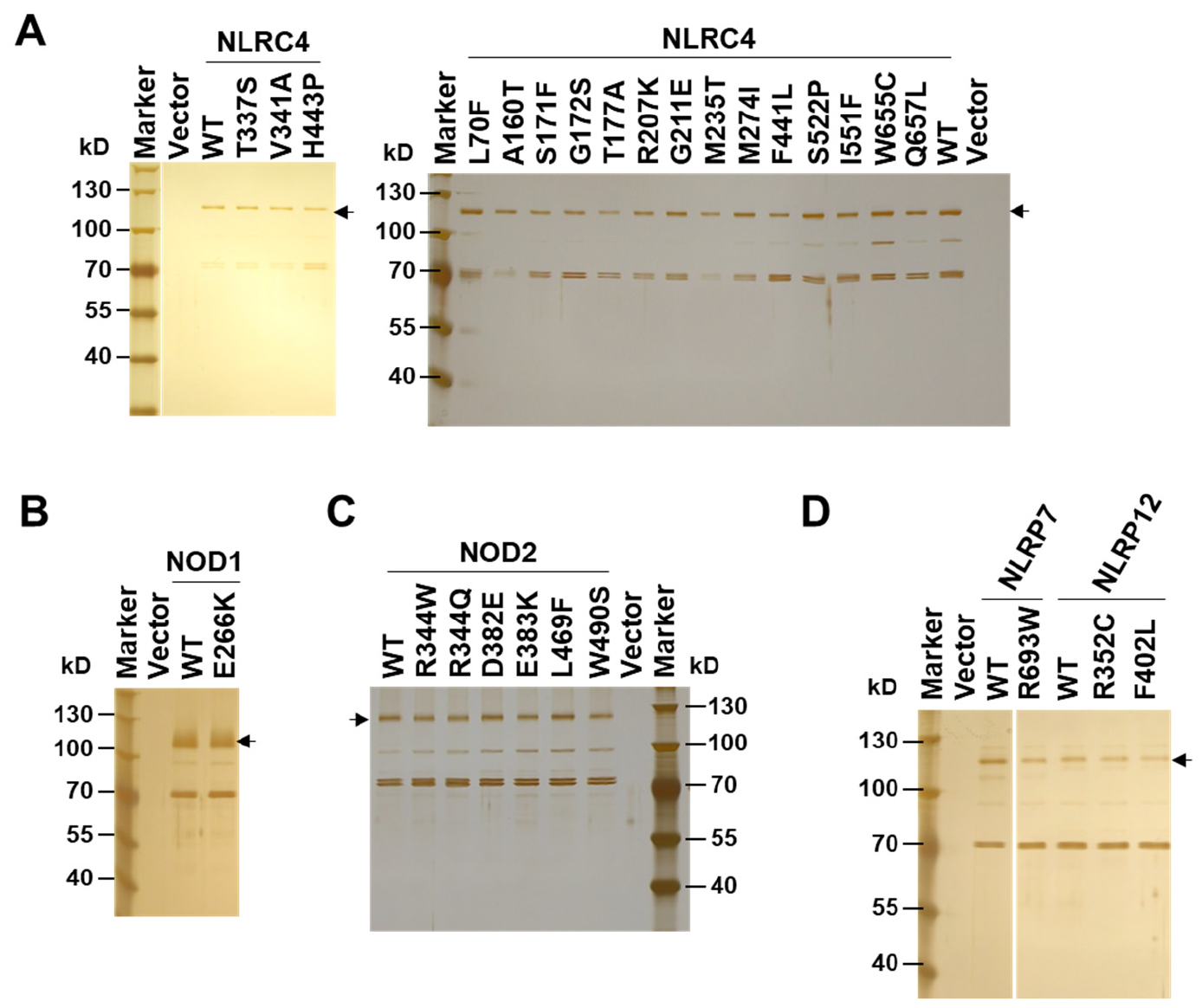
Silver staining of purified NLRC4, NOD1, NOD2, NLRP7 and NLRP12 proteins, related to Figure 2. **(A-D)** Representative image of silver staining of the purified Flag-tagged proteins as indicated by an arrow.

## STAR★METHODS

Detailed methods are provided in the online version of this paper and include the following:

- **KEY RESOURCES TABLE**
- **RESOURCE AVAILABILITY**
  - Lead contact
  - Materials availability
  - Data and code availability
- **EXPERIMENTAL MODEL AND SUBJECT DETAILS**
  - Cell lines
- **METHOD DETAILS**
  - Lentivirus production and infection
  - CRISPR-Cas9 knockout cells
  - Stable cell lines
  - Cell stimulation
  - Recombinant protein preparation
  - NLR proteins preparation
  - ATPase activity assay
  - Co-immunoprecipitation
- **QUANTIFICATION AND STATISTICAL ANALYSIS**

## KEY RESOURCES TABLE

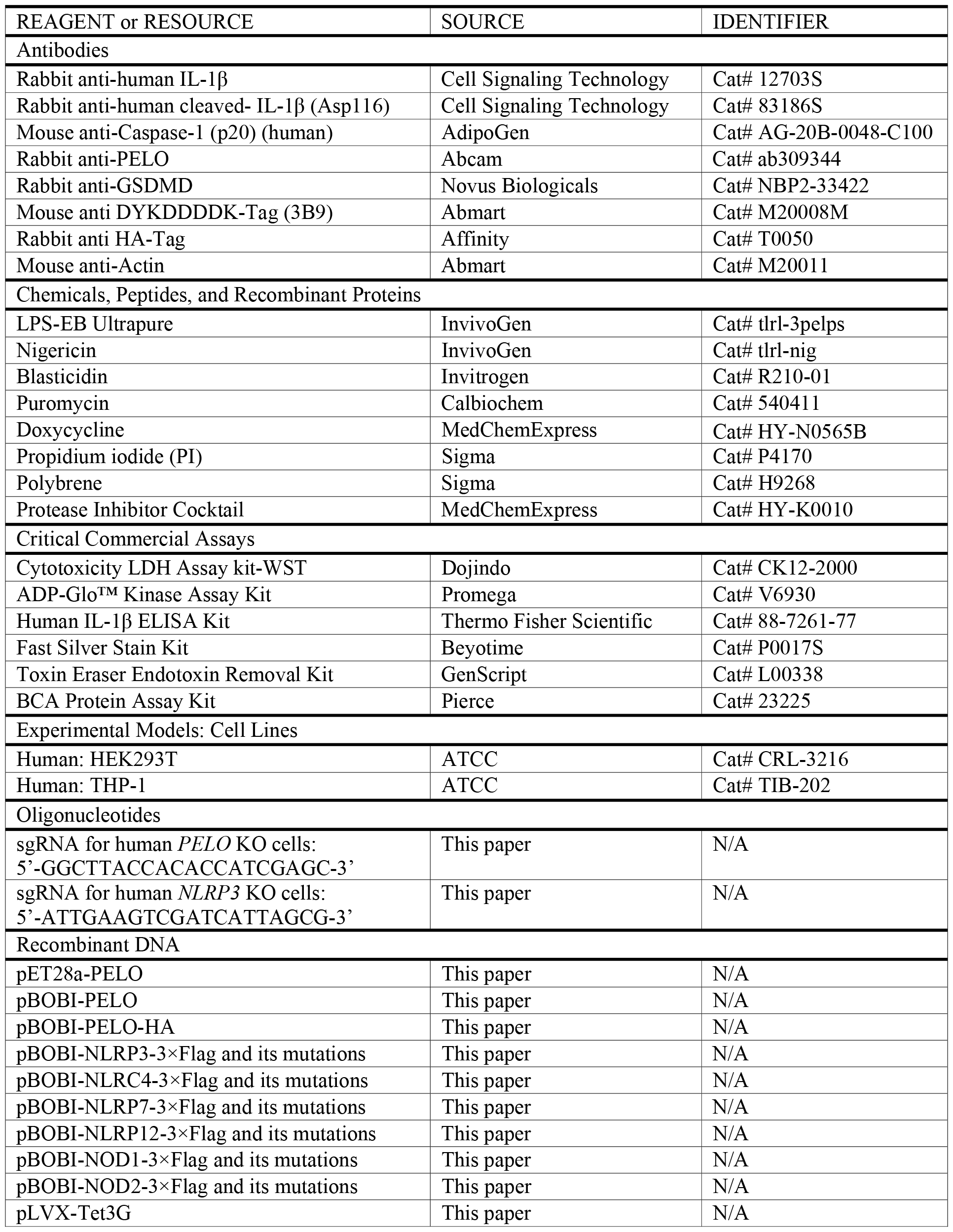
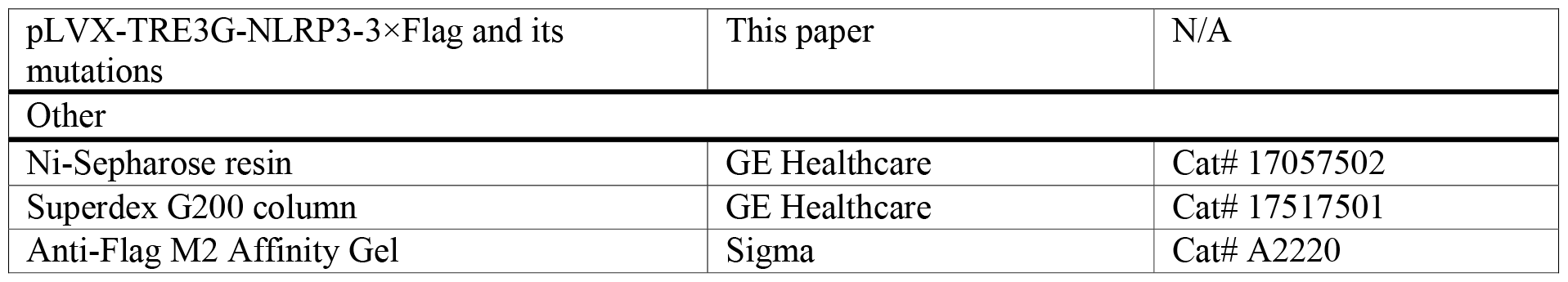

## RESOURCE AVAILABILTY

### Lead contact

Further information and requests for resources and reagents should be directed to and will be fulfilled by the Lead contact, Jiahuai Han.

### Materials availability

All plasmids, reagents, cell lines and mouse lines generated in this study are available from the Lead contact.

### Data and code availability

- Immunoblot images data and microscopy images reported in this paper will be shared by the Lead contact upon request.
- This paper does not report original code.
- Any additional information required to reanalyze the data reported in this paper is available from the Lead contact upon request.

## EXPERIMENTAL MODEL AND SUBJECT DETAILS

### Cell lines

The HEK293T cells were grown in DMEM supplemented with 10% FBS, and THP-1 cells were cultured in RPMI 1640 medium supplemented with 10% FBS. All cells were maintained at 37 °C in a 5% CO2 incubator.

## METHOD DETAILS

### Lentivirus production and infection

The lentiviral vectors, containing cDNAs or gRNAs of interest, were transfected into HEK293T cells along with lentivirus-packing plasmids (PMDL/REV/VSVG) using the calcium phosphate precipitation method. After 12 hours, the cell culture medium was changed and the virus-containing medium was collected 30 hours later. THP-1 cells were infected with lentiviral particles in the presence of 10 μg ml^−1^ polybrene and then centrifuged at 2,500 rpm for 60 min.

### CRISPR-Cas9 knockout cells

The target sequences used were 5 ‘-ATTGAAGTCGATCATTAGCG-3 ‘ for human *NLRP3* and 5 ‘-GGCTTACCACACCATCGAGC-3 ‘ for human *PELO*. To construct the knockout cell lines, gRNA was transduced into THP-1 cells via lentiviral delivery, followed by blasticidin (10 μg ml^−1^) selection. Single-cell clones were obtained through limiting dilution cloning in 96-well plates, and subsequently screened for the expression of the indicated genes using western blot. The selected knockout clones were further confirmed by DNA sequencing.

### Stable cell lines

The Tet-On system was used for inducible expression of NLRP3 mutations in THP-1 cells. Initially, THP-1 cells were infected with lentivirus encoding the Tet-On 3G transactivator protein and subsequently subjected to selection using blasticidin (10 μg ml^-1^). The resulting selected cells were utilized for subsequent experiments and further transduced with lentivirus carrying NLRP3 WT or mutations, followed by puromycin (10 μg ml^-1^) selection.

### Cell stimulation

THP-1 cells were initially primed with 1 mg ml^-1^ LPS for 4 hours in RPMI 1640 medium supplemented with 10% FBS. Following priming, the medium was replaced with serum-free RPMI 1640 medium containing 1 μg ml^-1^ doxycycline. Six hours later, the cells were stimulated with 5 μM nigericin for 1 hour. After stimulation, both culture supernatants and cell lysates were collected for immunoblotting analysis, with the cell supernatants were utilized for LDH assay (Cytotoxicity LDH Assay kit-WST, Dojindo Molecular Technologies) and ELISA analysis (human IL-1β ELISA Kit, Thermo Fisher Scientific) according to the manufacturer ‘s instructions.

### Recombinant protein preparation

The C-terminal 6×His-tagged PELO protein was expressed in *E. coli* BL21 (DE3) strain (Novagen, Merck) by overnight culturing at 30 °C in an auto-induction medium. The PELO protein was purified with Ni^2+^-NTA-agarose (Qiagen) and followed by further purification by Superdex 200 10/30 prepacked column (GE Healthcare). The protein concentration was determined using the BCA method with BSA as the standard.

### NLR proteins preparation

The NLR proteins were purified as previously described^5^. Briefly, *PELO* KO HEK293T cells were transiently transfected with plasmids encoding 3×Flag-tagged NLRs. 36 hours after transfection, the cells were washed twice in cold PBS and lysed in lysis buffer (50 mM HEPES, pH7.4, 150 mM NaCl, 1% NP-40) supplemented with Protease Inhibitor Cocktail. The lysates were incubated at 4°C on a rotation platform for 30 min and then centrifuged at 20, 000 g for 30 min at 4°C The supernatants were pre-cleaned with IgG-Agarose at 4°C on rotation for 2 hours, followed by incubation with Anti-Flag M2 Affinity Gel for 3 hours at 4°C on rotation. The beads were washed with wash buffer A (50 mM HEPES, pH7.4, 300 mM NaCl, 1% NP-40) for three times, followed by another three times of wash with wash buffer B (50 mM HEPES, pH7.4, 150 mM NaCl, 5% Glycerol, 10 mM MgCl_2_, 0.1% NP-40). The NLR proteins were finally eluted with wash buffer B containing 200 μg ml^-1^ 3×Flag peptide.

### ATPase activity assay

Assay was carried out using the ADP-Glo kinase assay kit in 384 Flat White Plates (142761, Thermo Fisher Scientific) as previously described^5^. Briefly, respective purified NLR proteins were incubated with BSA or PELO-His protein at 37°C for 30 min in the reaction buffer (50 mM HEPES, pH7.4, 150 mM NaCl, 5% Glycerol, 10 mM MgCl_2_, 0.1% NP-40, 1 mM DTT). Further, a standard containing 20 μM ATP, 16 μM ATP and 4 μM ADP, 8 μM ATP and 12 μM ADP, or 20 μM ADP was prepared in the same buffer. After incubation, ATP (20 μM in final) was added to the mixture and further incubated at 37°C for another 60 min. The reaction is stopped by addition of ADP-Glo reagent and further incubated for 40 min at room temperature. After the addition of kinase detection buffer, samples were incubated for another 40 min and luminescence read out using a Spark 20M microplate reader (Tecan) with an integration time of 1 s per well. For calculation of the individual values, a linear regression was calculated based on the ADP standard.

### Co-immunoprecipitation

HEK293T cells were lysed in NP-40 lysis buffer (25 mM HEPES, pH7.4, 150 mM NaCl, 1% NP-40) supplemented with Protease Inhibitor Cocktail. The lysates were incubated on a rocking platform for 30 min and then centrifuged at 20,000 g for 30 min at °C. Flag-tagged proteins were immunoprecipitated with Anti-Flag M2 Affinity Gel for 6 hours at °C. Beads containing protein complexes were washed with lysis buffer for three times. The immunocomplexes were eluted in SDS sample buffer and then analyzed by western blotting.

## QUANTIFICATION AND STATISTICAL ANALYSIS

No statistical methods were used to predetermine sample size. GraphPad Prism software was used for data analysis. Data are shown as mean ± standard deviation (SD).

